# In Rhesus monkeys, CSF-contacting neurons are the only neurons present in the medullo-spinal peri-ependymal zone

**DOI:** 10.1101/2023.03.29.534787

**Authors:** Anne Kastner, Nicolas Wanaverbecq

## Abstract

In spinal cord and medulla, ependymal cells are organized in a monolayer forming the central canal (cc). In rodents, this region, also known as a stem cell niche, was shown to contain cerebrospinal fluid-contacting neurons (CSF-cNs). These neurons are GABAergic and because of their chemo- and mechanosensory properties they would represent a novel sensory system intrinsic to the central nervous system. In primates, little is known about these neurons and more generally about the region around the cc. Here, using immunohistochemical approaches, we investigated the organization of the cc region and CSF-cN properties in *Macaca mulatta* Rhesus monkey. In contrast to rodent, we observe along the whole medullo-spinal axis a large zone around the cc delimited by long radial ependymal fibers that is enriched with astrocytes and microglia but largely devoid of neuronal elements except for CSF-cNs. These primate CSF-cNs share with rodent CSF-cNs similar morphological and phenotypical features with a largely immature profile. Our data suggest that they extend their axons in the longitudinal axis to form fiber bundles close to the cc and we further show that CSF-cNs receive GABAergic and serotoninergic synaptic contacts on their soma and dendrite. Taken together our results reveal in *Rh.* monkey a specific organization of the region around the cc potentially forming a buffer zone between CSF and parenchyma where CSF-cNs would play a crucial role in the detection of CSF signals and their transmission to the central nervous system, a role that would need to be further investigated.

## INTRODUCTION

Along the medullo-spinal axis, a monolayer of ependymal cells (EC) forms the central canal (cc) wall and separates cerebrospinal fluid (CSF) from parenchyma. EC selectively express vimentin (Vim) with markers of stem cells (nestin, Sox2, 3CB2 and FoxJ1) (Meletis et al., 2008; Hamilton et al., 2009; Marichal et al., 2012) and this region is therefore considered as a stem cell niche where reparatory and regenerative processes have been reported following lesion (Sabelström et al., 2013). Interestingly, close to and within the EC layer (ECL) neurons in contact with the CSF (CSF-cNs) are also found and exhibit a characteristic morphology and phenotype that is conserved in all vertebrates (Agduhr, 1922; Bruni and Reddy, 1987; LaMotte, 1987; Stoeckel et al., 2003; Vígh et al., 2004; Marichal et al., 2009; Djenoune et al., 2014; Orts-Del’Immagine et al., 2014) . These CSF-cNs are bipolar GABAergic cells, they extend a single large dendrite toward the cc ended with a protrusion or ‘bud’ in contact with the CSF and their axons regroup in the ventral region of the spinal cord (SC) to form large bilateral horizontal fiber bundles (Stoeckel et al., 2003; Djenoune et al., 2017; Jurčić et al., 2021; Gerstmann et al., 2022). CSF-cNs selectively express the polycystin kidney disease 2-like 1 (PKD2L1) ion channel, a member of the transient potential receptors (TRP) superfamily with sensory properties (Djenoune et al., 2014; Jalalvand et al., 2014; Orts-Del’Immagine et al., 2014; Petracca et al., 2016). In adult animals, they further exhibit an incomplete neuronal maturity state with a persistent expression of doublecortine (DCX) and the homeobox transcription factor Nkx6.1, two markers of immature neurons and neuroblasts, paralleled by a low expression of neuronal nuclear protein (NeuN), a marker of mature neurons (Sabourin et al., 2009; Kutna et al., 2013; Orts-Del’Immagine et al., 2014, 2017). At the functional level, CSF-cNs were shown to exhibit a spontaneous unitary current activity carried by PKD2L1 (Orts Del’Immagine et al., 2015; Jalalvand et al., 2016a; Sternberg et al., 2018; Orts-Del’Immagine et al., 2020) and modulated by changes in extracellular pH and osmolarity (chemoreception) (Orts Del’Immagine et al., 2015; Jalalvand et al., 2016a) as well as CSF flux and SC bending (mechanoreception) (Böhm et al., 2016; Jalalvand et al., 2016b; Sternberg et al., 2018; Orts-Del’Immagine et al., 2020). Therefore, CSF-cNs are suggested to represent a novel sensory system intrinsic to the central nervous system. This sensory function was confirmed in behavioral study in the zebrafish larva where CSF-cN selective activation lead to a state-dependent modulation of locomotor activity (Fidelin et al., 2015; Hubbard et al., 2016) and more recently in the mouse where CSF-cNs appear to participate in the modulation of postural motor control (Gerstmann et al., 2022; Nakamura et al., 2023).

In non-human primates only little information is available about the region around the spinal cc (Becker et al., 2018). In *Rh.* monkey, EC in the lateral part of the cc were shown to be multiciliated while in the ventral and dorsal region they are uni- or biciliated, express nestin and are capable of proliferating (Alfaro-Cervello et al., 2014). CSF-cNs were also observed in this model (LaMotte, 1987; Alfaro-Cervello et al., 2014; Djenoune et al., 2014) and were shown to express PKD2L1 and GABAergic markers (Djenoune et al., 2014) or the vasoactive intestinal peptide (VIP) (LaMotte, 1987). Nevertheless, the whole organization of these neurons and of the cc zone remains largely unknown. Therefore, to better understand in primates the whole organization of the cc region as well as CSF-cN properties and phenotype along the antero-posterior axis dedicated studies are necessary.

Here, we show that in *Macaca mulatas* or Rhesus monkey (*Rh.* monkey in the rest of the text), in contrast with rodents, the parenchyma is separated from the cc by a large hyponeuronal zone containing long radiating EC fibers and enriched with astrocytes and microglia forming a peri-ependymal zone (PEZ). This PEZ is observed along the whole medullo-spinal axis with conserved properties and the only neurons present in this area are CSF-cNs. Further, CSF-cNs exhibit similar morphological and phenotypical properties as in rodents. Their axons appear to form horizontal fiber bundles around the cc while GABAergic and serotoninergic, but not glutamatergic or cholinergic, synaptic contacts are observed on CSF-cN soma and dendrite. Taken together our results indicate the presence in *Rh.* monkey of a zone around the cc where astrocytes, microglia and EC fibers would act as a filtering/buffering zone for circulating bioactive signals between CSF and parenchyma, while CSF-cNs would play a crucial role in the detection of these signals and their transmission to the central nervous system, a role that would need to be further investigated.

## RESULTS

### 1. The region around the central canal forms a hyponeuronal astroglial zone

We first analyzed the organization of the region around the *Rh.* monkey cc at the cervical (C7/C8) SC level and look for the distribution of EC and their fibers, neuronal cell bodies, dendrites, and axons as well as astrocytes. Labelling against Vim indicates as expected the presence of EC only around the cc (Fig. 1A_1_-D_1_) with their cell body forming the cc wall or EC layer (ECL) (Fig. 1A_2_-D_2_). In agreement with previous work (Alfaro-Cervello et al., 2014), we observed that Vim positive fibers originating from the ECL extend radially within the white matter (WM) over long distance (∼100-150 µm on both side) (Apkarian and Hodge, 1989) to form a large zone around the cc (Fig. 1A_2_-D_2_), called peri-ependymal zone (PEZ) in the rest of the text. Immunolabelling against the neurofilament H (NFH, non-phosphorylated form; Fig. 1A_3_) or neurofilament 160 kDa (NF160, Fig. 1B_3_) reveals a strong immunoreactivity in the whole grey matter (GM) and a more diffused one in the white matter (WM; Fig. 1A_3_ and B_3_). Surprisingly, only rare neuronal elements are visible in the PEZ (Fig. 1A_3_-A_4_ for NFH and B_3_-B_4_ for NF160). The merge images on the right confirm this atypical organization and indicate a neuron-free or hyponeuronal PEZ around the cc (Fig. 1A_4_-B_4_). Note in B_3_, the presence of numerous NF160 positive puncta in the dorsal region of the SC corresponding to longitudinal axonal projection to or from supra-spinal structures. The absence of mature neuron in the PEZ is further illustrated by immunolabelling against NeuN indicating that neuronal cell bodies are only present outside the PEZ (Fig. 1C_3_-C_4_). Finally, we looked for the astrocyte distribution around and within the PEZ and our data indicate (Fig. 1D) a strong immunolabelling against the glial fibrillary acidic protein (GFAP) within the Vim immunoreactive PEZ (Fig. 1D_3_ and 1D_4_). Taken together our results show that in *Rh.* monkey the region around the cc is a hyponeural zone enriched with astrocytes and delimited by fibers radiating from EC cell bodies.

**Figure 1:**
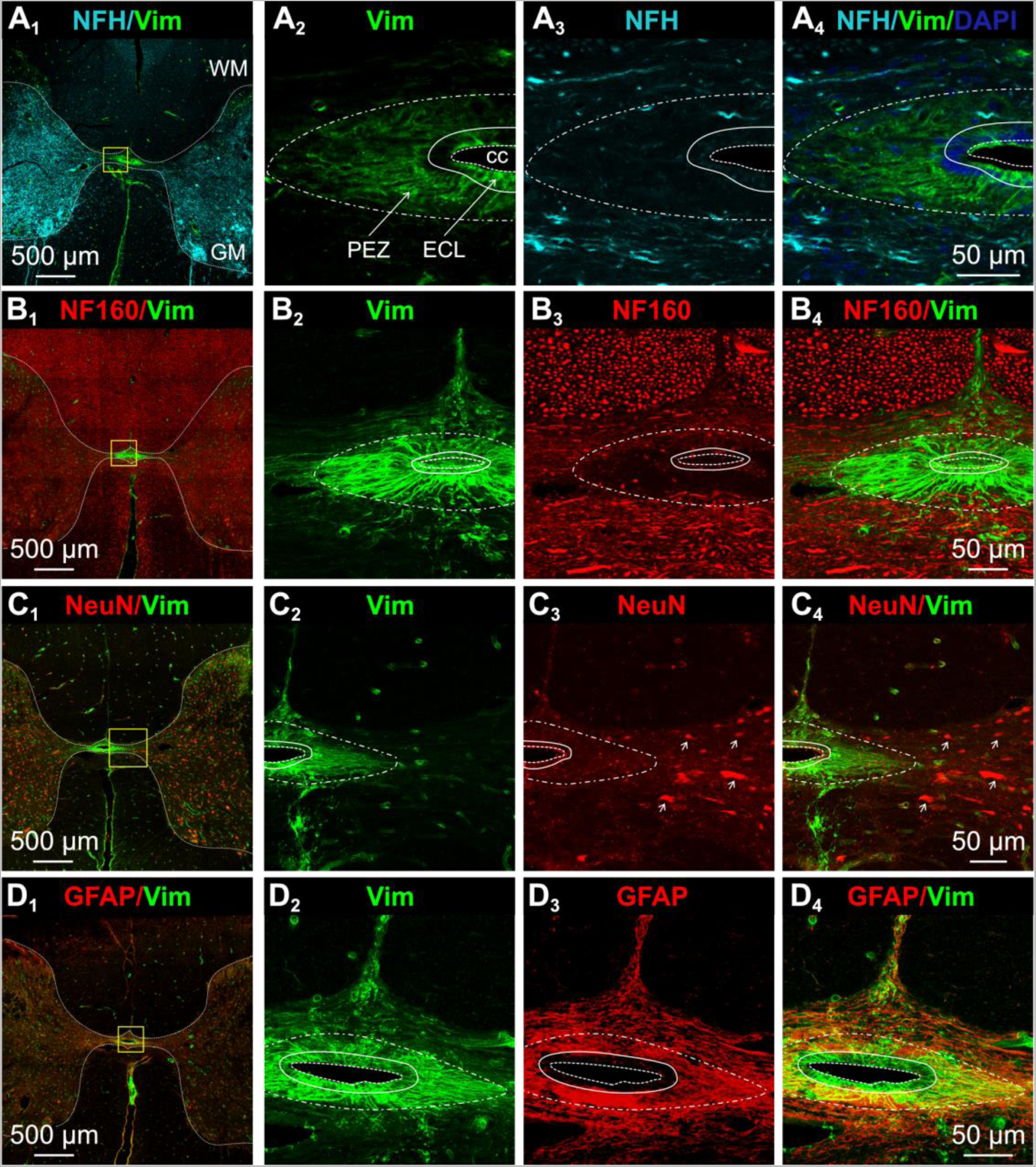
The region around the central canal is enriched with astrocytes and ependymal cell radiating fibers with only few neuronal elements. (**A1-D1**) Low magnification micrographs illustrating around the cc the expression of Vim, a marker for EC and fibers, along with neuronal markers (NFH, **A1** and NF160, **B1**), neuronal nucleus protein (NeuN, **C1**) and astrocytic markers (GFAP, **D1**) in transverse thin sections obtained from the caudal cervical segment of the *Rh.* monkey SC. (**A2-4**) High magnification images from **A1** for the region around the cc) indicating the presence of long radiating Vim positive fibers originating from soma in the ECL and extending to form the PEZ (**A2**). (**A3**) Immunolabelling against NFH to reveal neuronal elements around the cc. (**A4**) Merged image showing the distribution of Vim and NFH along with DAPI, a nuclear marker. Note the presence around the cc of the PEZ delineated by Vim+ radiating fibers and largely devoid of neuronal elements. (**B2-4**) High magnification images from **B1** showing Vim (**B2**) and NF160 (**B3**) labelling, respectively, to visualize EC and axons. The merged image in **B4** illustrates the organization of axonal fibers in the grey matter and around the PEZ. Note the absence of NF160 positive fibers within the PEZ as well as the presence of numerous puncta in the dorsal region of the white matter corresponding to longitudinal axonal projection. (**C2-4**) High magnification images from **C1** illustrating Vim (**C2**) and NeuN (**C3**, arrows in **C3** and **C4**), distribution around the cc. The merged image in **C4** confirms the absence of mature neurons within the PEZ. **D2-4**) High magnification images from **D1** showing the distribution of Vim (**D2**) and GFAP, an astrocytic marker (**D3**). The merged image in D4 indicates the presence of a region around the cc that is delineated by Vim positive fibers and enriched with astrocytes. Sections are oriented with the dorsal region on top and the regions illustrated in high magnification images corresponds to those indicated with yellow boxes in the low magnification images on the left. Color code for the labels on top of images corresponds to the illustrated markers. Dashed lines in **A1** to **D1** delineates the GM from the WM with the dorsal and ventral horns. Scale bars are shown in the images left and right. Dashed lines: cc; Full lines: ECL; Doted-dashed lines: PEZ.

### 2. CSF-contacting neurons are the only neurons present within the Peri-ependymal zone

In previous studies in mice, we reported that CSF-cNs do not present a high expression level for NeuN and that classical neuronal maturity markers are absent, while microtubule associated protein type 2 (MAP2) immunolabelling prove to be a very efficient marker to visualize these neurons in medullo-spinal tissues (Orts-Del’Immagine et al., 2014, 2017; Jurčić et al., 2021). We therefore performed a set of experiments using antibodies against MAP2 to look for CSF-cNs distribution and morphology around the *Rh.* monkey cc. In transverse SC sections at the low cervical level (C7/C8), we observed a dense MAP2 immunolabelling present in the whole GM that surround a region highly immunoreactive against Vim corresponding to the PEZ (Fig. 2A). However, in contrast to immunolabelling against NFH or NF160, MAP2 immunoreactivity is also observed within the PEZ that selectively and exclusively labels CSF-cNs (Fig. 2A_2_). These MAP2 positive CSF-cNs present the characteristic morphology with a small round soma (arrows) from which a large dendrite (arrowheads) projects to the cc ending with a protrusion in the lumen (asterisks). Although, in *Rh.* monkey CSF-cN morphology is similar to that observed in rodents (LaMotte, 1987; Djenoune et al., 2014; Orts-Del’Immagine et al., 2014; Jurčić et al., 2021), they are present within the ECL, beneath it or can be localized further away from the cc with dendrite extending over longer distances (> 50 µm) (Fig. 2A_2_, see also Fig. 4 and (5). Finally, CSF-cN cell bodies are surrounded by Vim positive radial fibers and their dendrites project to the cc along these fibers (Fig. 2A_3_). We further performed double immunolabelling analyses against NFH and MAP2 or β3-tubuline (Tuj1; β3Tub), a marker against cytoskeletal microtubules in the somato-dendritic and axonal compartments and confirm that the PEZ represents an hyponeural region where only CSF-cNs are present (Figure 2B_1_) and indicate that β3-Tub also allows to selectively visualized CSF-cNs within the PEZ (Fig. 2B_2_). Next, we looked for the microglia distribution within the SC tissue and reveal a strong immunoreactivity against the ionized calcium-binding adaptor molecule 1 (Iba1), a selective marker for microglia (Sasaki et al., 2001), in the parenchyma (Fig. 2C). Interestingly, we also observe dense Iba1 immunoreactivity within the PEZ and around the cc where Iba1 positive fibers can be visualize crossing the ECL towards the lumen (arrows in Fig. 2C) or close to MAP2 positive CSF-cN fibers. The PEZ is therefore a region delimited by Vim positive radial fibers enriched with astrocyte and microglia where the only neurons present are CSF-cNs. Analysis of longitudinal sections further confirms this unique PEZ organization delimited on its edge by the parenchyma (strong immunoreactivity against MAP2; Fig. 2D_1_), while MAP2 positive CSF-cNs are distributed at high density within the PEZ along the cc with their dendrite projecting perpendicular to the longitudinal axis (Fig. 2D_1_) and parallel to Vim positive fibers originating from EC cell bodies (Fig. 2D_2_). Next, in agreement with earlier reports in fish and mouse (Djenoune et al., 2014; Orts-Del’Immagine et al., 2014; Jurčić et al., 2021), CSF-cNs selectively express PKD2L1 (although PKD2L1 immunoreactivity is low) (Fig. 2E) and are GABAergic (GAD67, Fig. 2F). To confirm whether the organization observed in *Rh.* monkey for the region around the cc is specific to this animal model, we repeated a set of experiments in SC sections obtained from mice and rats, and in contrast to the data obtained in *Rh.* monkey, we did not observe a specific PEZ in rodent tissues. The cc appears rather directly in contact with both astrocytes (GFAP, Fig. 3A_1_ and 3B_1_) and neuronal elements (NFH, Fig. 3A_2_ and 3B_2_). Further, in rodent Vim positive radial fibers extend from the ECL within MAP2 positive elements in the parenchyma (Vim and MAP2, Fig. 3C_1_ and 3D_1_) and surround PKD2L1 positive CSF-cNs (Fig. 3E_2_ and 3F_2_). Finally, immunolabelling against NeuN in rodent tissues reveals the presence of neurons contiguous to the cc (Fig. 3C_2_ and 3D_2_). Altogether, these results indicate that, in *Rh.* monkey, CSF-cNs exhibit similar morphological and phenotypical features than in rodent. However, in this primate model, the region around the cc in characterize by a unique PEZ where CSF-cNs are the only neurons present.

**Figure 2:**
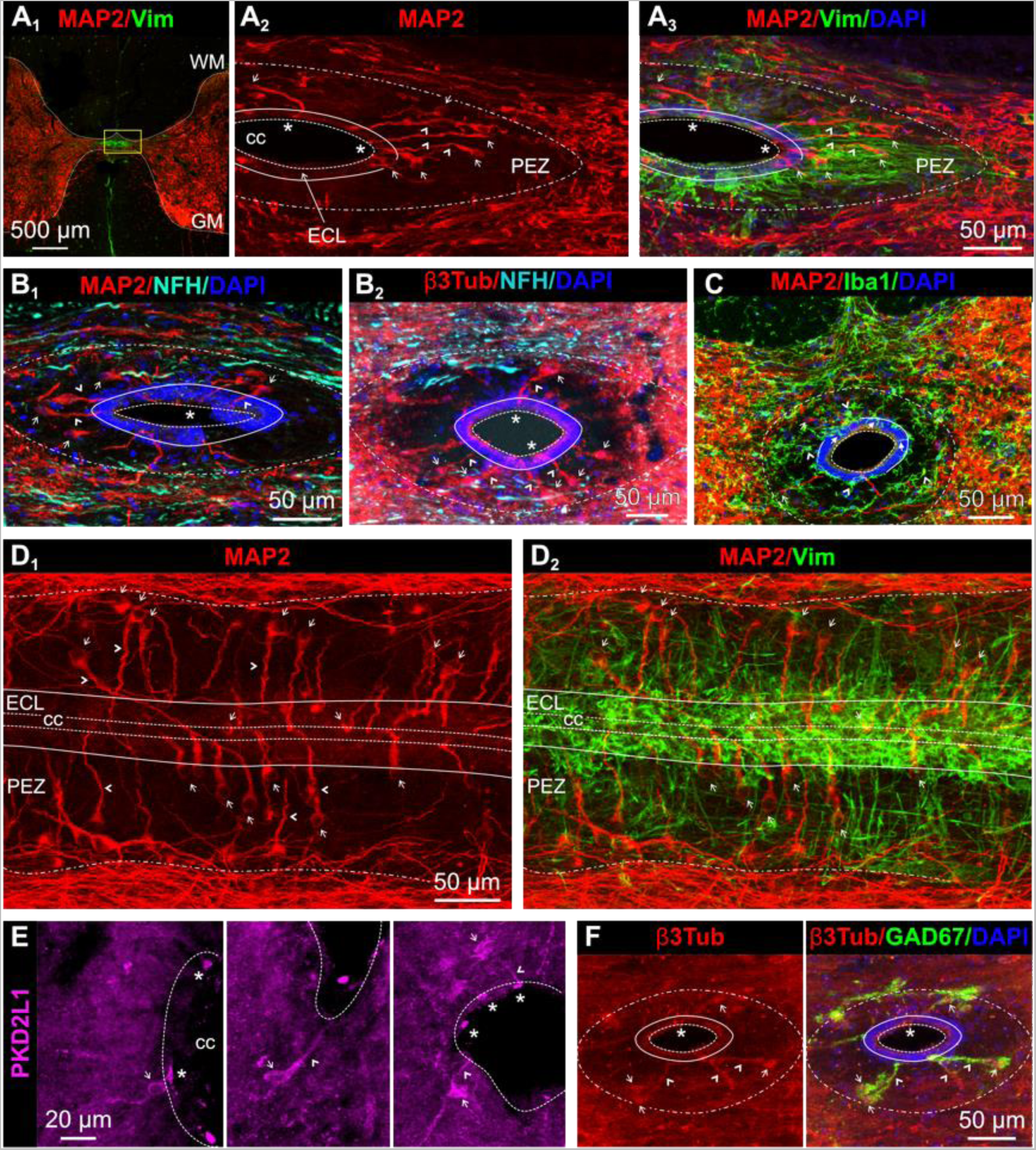
The peri-ependymal zone contains CSF-contacting neurons, the only neurons present in this region. (**A1**) Low magnification micrographs illustrating the distribution of Vim, to reveal EC and fibers, and MAP2, a marker for the somato-dendritic compartments, in one transverse thin section obtained from the caudal cervical segment of the SCSC. Note that MAP2 is intensively expressed in the whole GM. (**A2**) MAP2 immunolabelling reveals the presence of neurons in the PEZ. These neurons exhibit the characteristic morphology of CSF-contacting neurons (CSF-cNs), with a small soma (arrows) localized around the cc and projecting a single large dendrite (arrowheads) ended with a round protrusion or bud (asterisks) in the cc lumen. Note also the intense MAP2 immunolabelling outside the PEZ, in the GM. The merged image in **A3** shows that CSF-cNs (MAP2) are the only neurons present in the PEZ and extend their dendrites along the Vim positive fibers. DAPI: nuclear marker. **(B)** MAP2/NFH co-immunolabelling (**B1**) confirming that CSF-cNs are restricted to the NFH negative hyponeuronal PEZ. **(B2)** β3Tub, a marker for somato-dendritic and axonal compartments, along with NFH immunolabelling allows CSF-cN visualization in the PEZ devoid of NFH immunoreactivity. **(C)** Co-immunolabelling against MAP2 and Iba1, a marker for microglial cells, illustrating their presence in the parenchyma, the PEZ (arrowheads) close to CSF-cNs (arrows) as well as around the cc with projection (filled arrows) within the ECL. (**D**) Horizontal thin section obtained at the level of the cc (dashed lines) of a cervical SCSC section and immunolabelled against MAP2 (**D1 and 2**) and Vim (**D2**) to reveal CSF-cNs and EC or fibers, respectively. Note that MAP2 immunolabelling indicates the distribution in the antero-posterior axis of CSF-cNs within the PEZ along Vim positive fibers. (**E**) Micrographs illustrating PKD2L1 selective expression in the soma (arrows), dendrite (arrowheads), and bud (asterisk) of CSF-cNs in transverse thin sections prepared from cervical SCSC. **F**) Transverse thin section (caudal cervical level) immunolabelled against Tub (Left) and GAD67 (Right), a selective marker of GABAergic neurons. For abbreviations see Figure 1. The region illustrated in the high magnification images in **A2** and **A3** corresponds to that indicated with a yellow box in **A1** (left). Color code for the labels on top of images corresponds to the illustrated markers. Scale bars are shown on the images.

**Figure 3:**
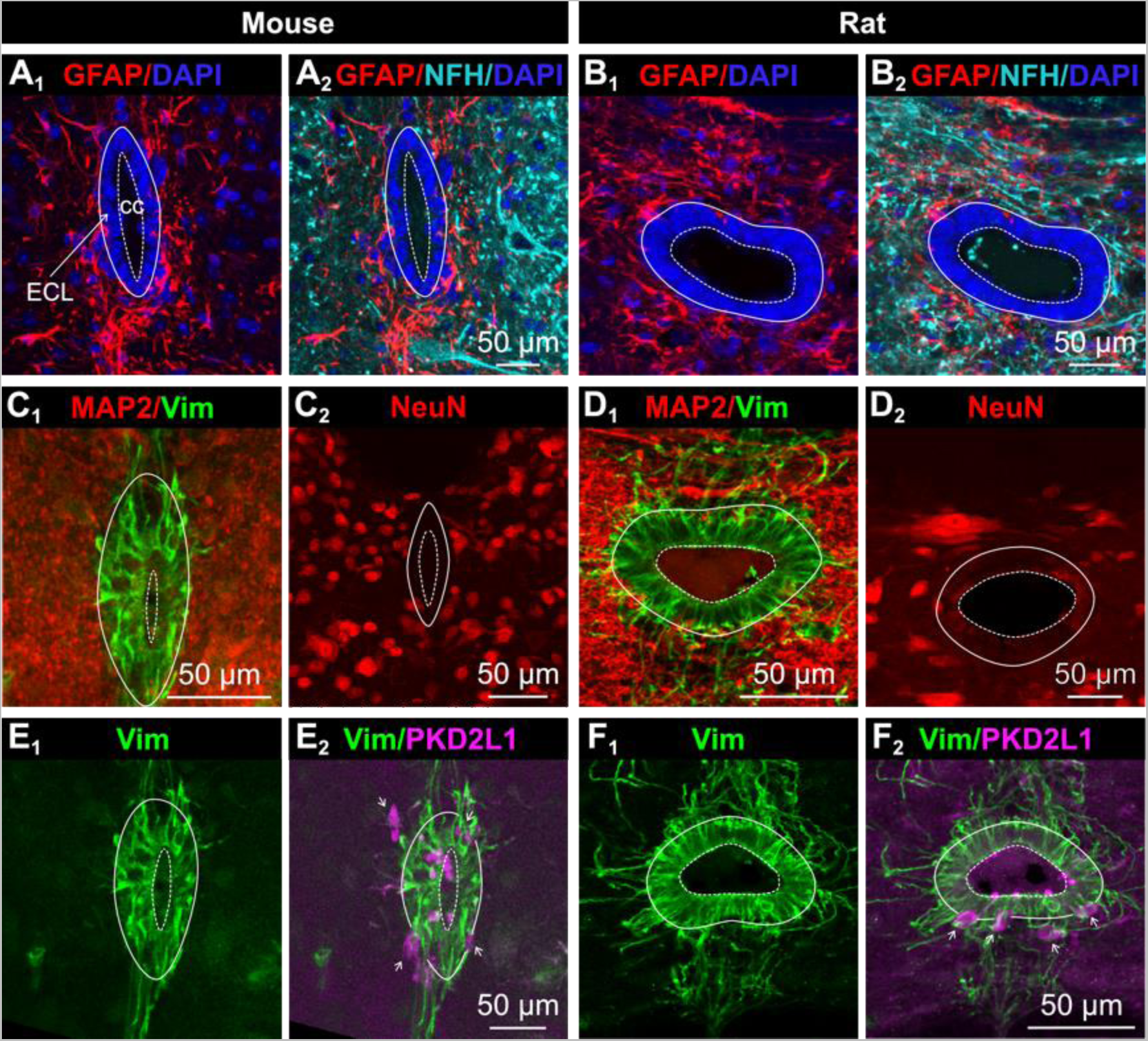
In rodents the ependymal cell layer is contiguous to the parenchyma. Organization of the region around the cc in cervical sections obtained from mouse (**A, C** and **E**) and rat (**B, D** and **F**) SC. (**A** and **B**) Images of the astrocyte distribution (GFAP; **A1** and **B1**) along with nuclear labeling (DAPI). (**A2** and **B2**) Merged micrographs showing the distribution for the immunolabelling against GFAP and NFH, a neuronal marker. (**C** and **D**) Images of the immunolabelling against Vim and MAP2 (**C1** and **D1**) and against NeuN (**C2** and **D2**) around the cc. (**E** and **F**) Images of the immunolabelling against Vim (**E1** and **F1**) and merged images of Vim and PKD2L1 staining (**E2** and **F2**). The images confirm the absence of a PEZ in rodent illustrated by the close apposition of MAP2 and Vim labelling and the presence of NeuN immunolabelling close to the cc (compare with Figures 1 and 2). Note that CSF-cNs selectively express PKD2L1 and are present within or close to the ECL in direct contact with the parenchyma. For abbreviations see Figure 1. Color code for the labels on top of images corresponds to the illustrated markers. Scale bars are shown on the images. Arrows: CSF-cNs.

### 3. PEZ and CSF-cN organization and properties along the rostro-caudal axis of the central canal

According to previous reports, the whole SC size and GM shape vary along the different rostro-caudal levels (Apkarian and Hodge, 1989) and this observation is confirmed by our results (Fig. 4). To determine whether the unique PEZ organization observed at the cervical level is conserved along the whole cc antero-posterior axis, we performed histological and quantitative analysis of the regions around the cc from the medulla to the sacral segments (Fig. 4A_1_ to 4E_1_). Our results indicate that the PEZ anatomy is conserved at all medullo-spinal segments and that neuronal elements identified with immunolabelling against NFH surround a large zone around the cc that is selectively enriched with astrocytes (GFAP, Fig. 4A_2_ to E_2_). When considering the PEZ and cc orientation and shape in the segments of interest, we noted that both extended in the dorso-ventral axis at the medulla, mid-cervical as well as in the lumbar and sacral segments (Fig. 4). In contrast, at the high-cervical (Fig. 4B), low-cervical (Fig. 1) and thoracic (Fig. 4C) levels, the PEZ and cc are rather oriented in the horizontal axis. We next looked for the presence of CSF-cNs in the PEZ in the same segments using staining against MAP2, and our data confirm that CSF-cNs are present in the PEZ along the whole rostro-caudal axis of the cc but more importantly that they are the only neuronal population in these regions (Fig. 4A_3_-E_3_). When looking more closely at CSF-cN distribution in the different segments, our results suggest a differential organization. At the medullar and cervical levels, CSF-cNs are mainly found within the PEZ (Fig. 4A_3_ and 4B_4_) at various distance from the ECL, while from thoracic to sacral segments, CSF-cNs are mostly found near or within the ECL or in a dorso-ventral position (Fig. 4C_3_ to 4E_3_). Immunolabelling against β3-Tub confirms this observation and further indicates the presence of axonal structures originating from CSF-cNs (4A_4_ to 4E_4_). To further characterize the organization of the PEZ in our study, we performed a quantitative analysis for the PEZ and cc anatomy as well as of CSF-cN distribution (Fig. 5 and Table 1). First, we measured from the medullar to the sacral levels the PEZ width and we indicate that it is the largest (around 80 µm) at the medullar and caudal cervical levels (Med and cCerv; Fig. 5A and Table 1A). The PEZ width is smaller at the rostral and mid-cervical (around 50 µm; rCerv and mCerv) and further decreases at more caudal segments to be around 30 µm (from thoracic to sacral segments). Finally, CSF-cN cell bodies are surrounded by Vim positive radial fibers and their dendrites project to the cc along these fibers (Fig. 2A_3_). We further performed double immunolabelling analyses against NFH and MAP2 or β3-tubuline (Tuj1; β3Tub), a marker against cytoskeletal microtubules in the somato-dendritic and axonal compartments and confirm that the PEZ represents an hyponeural region where only CSF-cNs are present (Figure 2B_1_) and indicate that β3-Tub also allows to selectively visualized CSF-cNs within the PEZ (Fig. 2B_2_). Next, we looked for the microglia distribution within the SC tissue and reveal a strong immunoreactivity against the ionized calcium-binding adaptor molecule 1 (Iba1), a selective marker for microglia (Sasaki et al., 2001), in the parenchyma (Fig. 2C). Interestingly, we also observe dense Iba1 immunoreactivity within the PEZ and around the cc where Iba1 positive fibers can be visualize crossing the ECL towards the lumen (arrows in Fig. 2C) or close to MAP2 positive CSF-cN fibers. The PEZ is therefore a region delimited by Vim positive radial fibers enriched with astrocyte and microglia where the only neurons present are CSF-cNs. Analysis of longitudinal sections further confirms this unique PEZ organization delimited on its edge by the parenchyma (strong immunoreactivity against MAP2; Fig. 2D_1_), while MAP2 positive CSF-cNs are distributed at high density within the PEZ along the cc with their dendrite projecting perpendicular to the longitudinal axis (Fig. 2D_1_) and parallel to Vim positive fibers originating from EC cell bodies (Fig. 2D_2_). Next, in agreement with earlier reports in fish and mouse (Djenoune et al., 2014; Orts-Del’Immagine et al., 2014; Jurčić et al., 2021), CSF-cNs selectively express PKD2L1 (although PKD2L1 immunoreactivity is low) (Fig. 2E) and are GABAergic (GAD67, Fig. 2F). To confirm whether the organization observed in *Rh.* monkey for the region around the cc is specific to this animal model, we repeated a set of experiments in SC sections obtained from mice and rats, and in contrast to the data obtained in *Rh.* monkey, we did not observe a specific PEZ in rodent tissues. The cc appears rather directly in contact with both astrocytes (GFAP, Fig. 3A_1_ and 3B_1_) and neuronal elements (NFH, Fig. 3A_2_ and 3B_2_). Further, in rodent Vim positive radial fibers extend from the ECL within MAP2 positive elements in the parenchyma (Vim and MAP2, Fig. 3C_1_ and 3D_1_) and surround PKD2L1 positive CSF-cNs (Fig. 3E_2_ and 3F_2_). Finally, immunolabelling against NeuN in rodent tissues reveals the presence of neurons contiguous to the cc (Fig. 3C_2_ and 3D_2_). Altogether, these results indicate that, in *Rh.* monkey, CSF-cNs exhibit similar morphological and phenotypical features than in rodent. However, in this primate model, the region around the cc in characterize by a unique PEZ where CSF-cNs are the only neurons present. Next, we analyzed the cc area (Fig. 5B) at the same levels of interest and found the higher values at the medulla (Med) and sacral (Sac) levels (around 7 x 10^3^ µm^2^). In more caudal regions between the rostral cervical and lumbar segments, the cc area ranges from 3 to 5 x 10^3^ µm^2^ with no significant differences ((Figure 5B and Table 1B; see Figure legend for details). We further analyzed the CSF-cN density ((Figure 5C and Table 1C; number of CSF-cNs/10 µm tissue depth) along the rostro-caudal axis of the cc and found the largest cell density at the mid and caudal cervical levels ((Figure 5C; cell number between 10 and 15) compared to the other segments where the density is on average around 8 CSF-cNs ((Figure 5C and Table 1C; see Figure legend for details). Finally, regarding CSF-cN distance from the cc (*i.e.* the length of their dendrite), our measurements indicate that CSF-cN cell bodies are on average at 50 µm from the cc in the medulla and cervical regions, *i.e.* the most rostral zones, except at the mid-cervical level where CSF-cNs are significantly closer ((Figure 5D and Table 1). In more caudal segments, CSF-cNs are found closer to the cc although with a larger variability and an average distance around 20 to 25 µm ((Figure 5D and Table 1). In the more anterior segments (Med and Cerv), CSF-cNs could be found at distance as high as 100 µm right at the edge of the PEZ (see Fig. 2A_2_ and 2B_1_). However, at all levels, CSF-cNs are distributed in the whole PEZ as well as within the ECL (intra-ependymal). Note that due to the maker used (MAP2) to visualize CSF-cNs and perform the quantification, one cannot exclude the presence of CSF-cNs within the parenchyma. Altogether, our results indicate that the PEZ and cc anatomy varies along the rostro-caudal axis with the widest PEZ and larger cc observed mostly in the anterior segments. This anatomy is accompanied by CSF-cNs at a higher density and larger distance to the cc for the same anterior segments.

**Figure 4:**
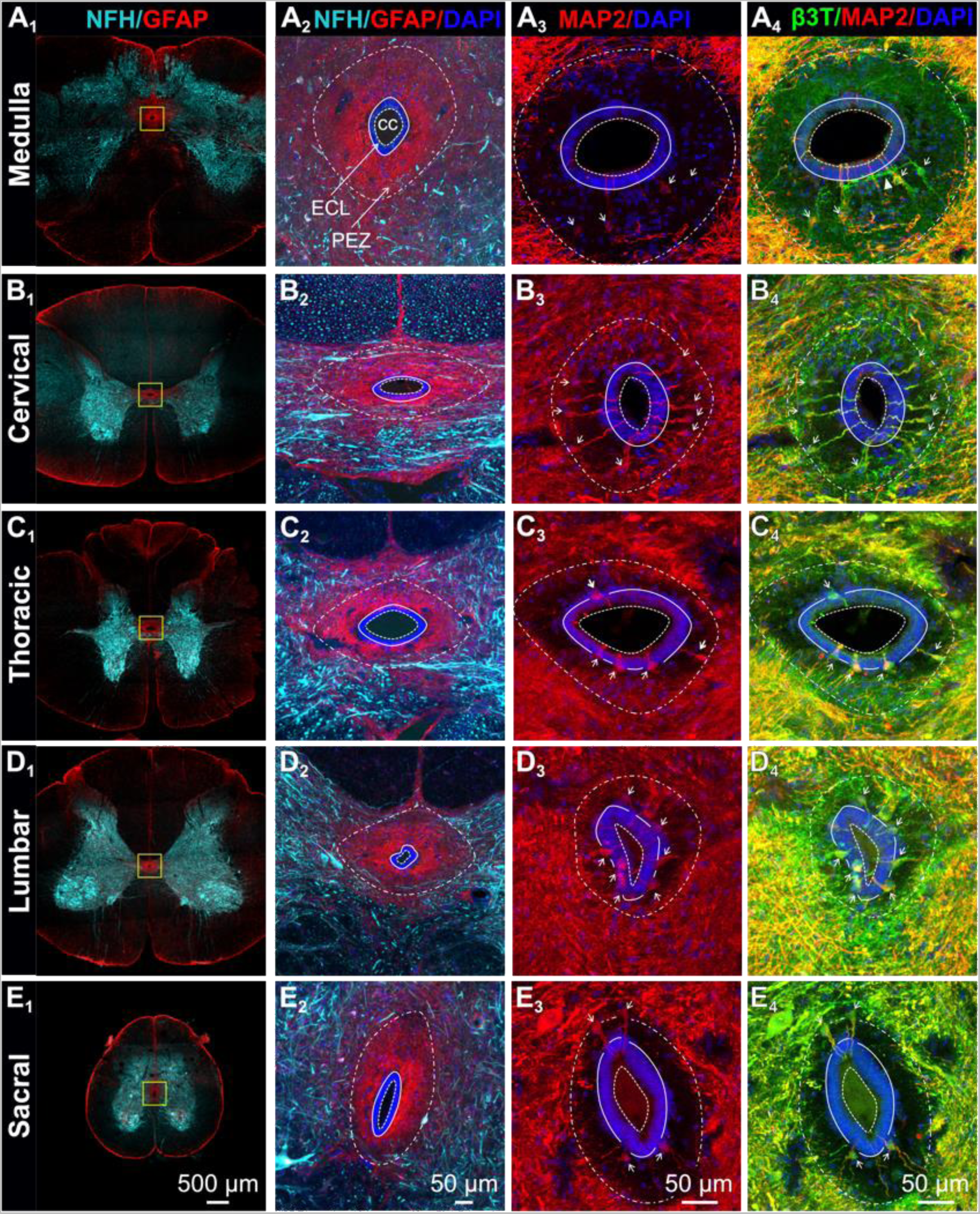
In macaque, the peri-ependymal zone organization is conserved along the whole rostro-caudal axis of the central canal. **(A1**-**E1**) Micrographs at low magnification illustrating the anatomy of the medulla and the SC (cervical to sacral) along the antero-posterior axis in transverse sections with the characteristic organization of the GM at each level. (**A2**-**E2**) High magnification images of the region around the cc from the sections illustrated from **A1** to **E1** (yellow squares) and showing the immunolabelling against NFH, GFAP and DAPI. (**A3**-**E3**) Representative images for the PEZ organization with the presence of CSF-cNs around the cc and labelled with MAP2 and DAPI. (**A4**-**E4**) Images showing the co-immunolabelling against β3Tub and MAP2 to reveal axonal (β3Tub only; arrowhead) as well as somato-dendritic compartments β3Tub and MAP2). For abbreviations see Figure 1. Arrows point to CSF-cNs. Color code for the labels on top of images corresponds to the illustrated markers. Scale bars are shown on the images.

**Figure 5:**
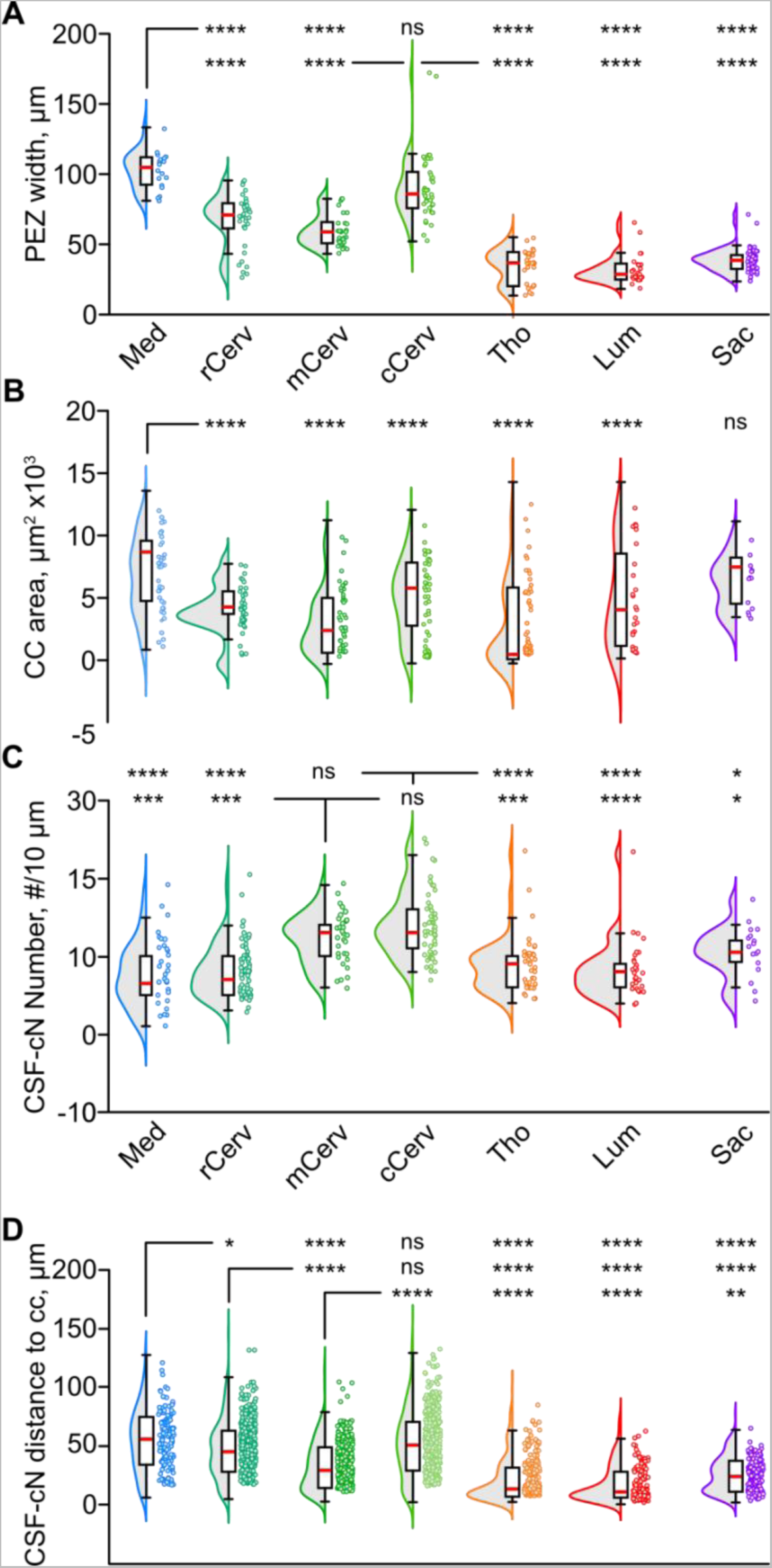
Properties of the peri-ependymal zone and CSF-cNs distribution along the medullo-spinal axis. Summary graphs for the quantification of the PEZ width (**A**), the cc lumen area (µm^2^ x 10^3^, **B**), the number of CSF-cNs (**C**) and their distance to the cc (**D**) at the medulla (Med), the rostral, mid, and caudal cervical zone (rCerv, mCerv and cCerv), the thoracic (Tho), the lumbar (Lum) and the sacral (Sac) segments. For each graph, whiskers boxplots are presented with superimposed violin plots for the data point density and distribution. Statistical significance was analyzed using a non-linear multi-effect test: (**A**) χ^2^ = 621.84, df = 6 and p(χ^2^) < 2.2 x 10^-16^;(**B**) χ^2^ = 56,11, df = 6 and p(χ^2^) = 2.763 x 10^-10^; (**C**) χ^2^ = 113.36, df = 6 and p(χ^2^) < 2.2 x 10^-16^; (**D**) χ^2^ = 416.11, df = 6 and p(χ^2^) < 2.2 x 10^-16^. Pairwise post-hoc comparison using Sidak method: *, < 0.05; **, < 0.01; ***, < 0.001 and ****<0.0001 (see detail in Table 1).

**Table 1:**
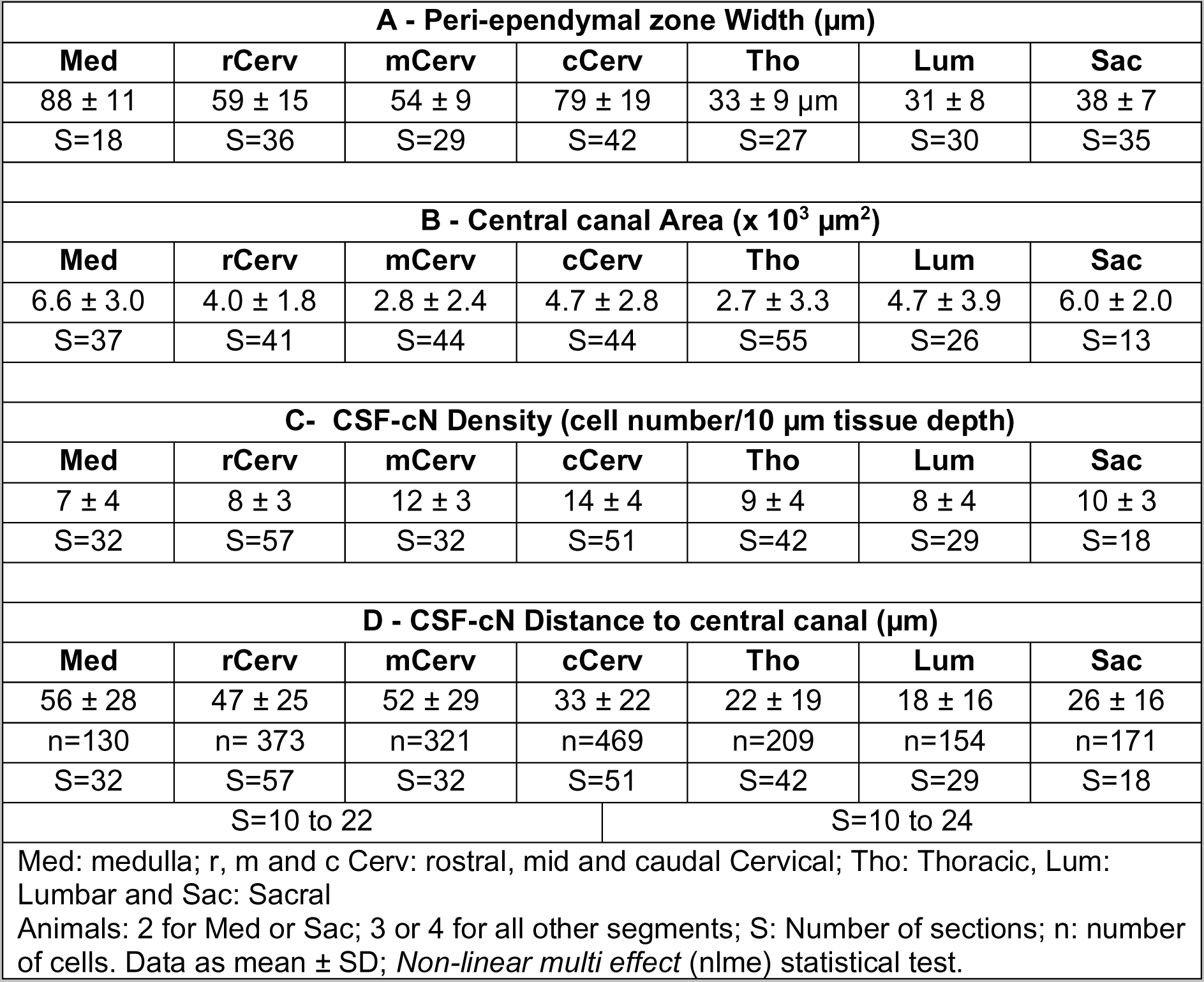
Quantitative analysis of the Peri-ependymal Zone and CSF-cNs properties

| A - Peri-ependymal zone Width (µm) |  |  |  |  |  |  |
| --- | --- | --- | --- | --- | --- | --- |
| Med | rCerv | mCerv | cCerv | Tho | Lum | Sac |
| 88 ± 11 | 59 ± 15 | 54 ± 9 | 79 ± 19 | 33 ± 9 µm | 31 ± 8 | 38 ± 7 |
| S=18 | S=36 | S=29 | S=42 | S=27 | S=30 | S=35 |
| B - Central canal Area (x 10<sup>3</sup> µm<sup>2</sup>) |  |  |  |  |  |  |
| Med | rCerv | mCerv | cCerv | Tho | Lum | Sac |
| 6.6 ± 3.0 | 4.0 ± 1.8 | 2.8 ± 2.4 | 4.7 ± 2.8 | 2.7 ± 3.3 | 4.7 ± 3.9 | 6.0 ± 2.0 |
| S=37 | S=41 | S=44 | S=44 | S=55 | S=26 | S=13 |
| C- CSF-cN Density (cell number/10 µm tissue depth) |  |  |  |  |  |  |
| Med | rCerv | mCerv | cCerv | Tho | Lum | Sac |
| 7 ± 4 | 8 ± 3 | 12 ± 3 | 14 ± 4 | 9 ± 4 | 8 ± 4 | 10 ± 3 |
| S=32 | S=57 | S=32 | S=51 | S=42 | S=29 | S=18 |
| D - CSF-cN Distance to central canal (µm) |  |  |  |  |  |  |
| Med | rCerv | mCerv | cCerv | Tho | Lum | Sac |
| 56 ± 28 | 47 ± 25 | 52 ± 29 | 33 ± 22 | 22 ± 19 | 18 ± 16 | 26 ± 16 |
| n=130 | n= 373 | n=321 | n=469 | n=209 | n=154 | n=171 |
| S=32 | S=57 | S=32 | S=51 | S=42 | S=29 | S=18 |
| S=10 to 22 |  |  | S=10 to 24 |  |  |  |
| Med: medulla; r, m and c Cerv: rostral, mid and caudal Cervical; Tho: Thoracic, Lum: Lumbar and Sac: Sacral |  |  |  |  |  |  |
| Animals: 2 for Med or Sac; 3 or 4 for all other segments; S: Number of sections; n: number of cells. Data as mean ± SD; *Non-linear multi effect* (nlme) statistical test. |  |  |  |  |  |  |

### 4. In *Rh.* monkey, CSF-cNs present a similar phenotype as in rodents and exhibit an intermediate neuronal maturity

In rodents, CSF-cNs are known to exhibit a low/intermediate neuronal maturity state (Orts-Del’Immagine et al., 2014, 2017; Petracca et al., 2016; Jurčić et al., 2021) characterized by the absence or low levels of NeuN (a marker for mature neurons) and doublecortin (DCX, a marker of immature neurons) along with Nkx6.1 expression (a homeobox transcription factor expressed in neuronal cells at early developmental stages and only conserved in some neural progenitors of the ventral spinal neuroepithelium in adult animals) (Sabourin et al., 2009; Kutna et al., 2013; Orts-Del’Immagine et al., 2014, 2017). To date, little is known about CSF-cN phenotype and maturity state in *Rh.* monkey. We reported above the absence of NeuN immunolabelling within the PEZ (see Fig. 1C) but to further explore CSF-cN maturity state in *Rh.* monkey, we performed a set of immunohistofluorescence experiments using classical maturity/immaturity markers. Here, we carried out double immunostaining against MAP2 (to visualize CSF-cNs) and NeuN to assess its level of expression selectively in CSF-cNs. Our results confirm the absence of NeuN immunoreactivity within the PEZ (Fig. 6A_1_) and further indicate that CSF-cNs in do not express NeuN (Fig. 6A_2_ and see inset). We next looked for DCX (Fig. 6B) and Nkx6.1 (Fig. 6C) expression in CSF-cNs and our data reveal that CSF-cNs do express both immaturity markers. The selective Nkx6.1 expression in CSF-cNs is further confirmed by the co-immunolabelling against PKD2L1 and Nkx6.1 observed in this neuronal population (Fig. 6D). The immature phenotype for CSF-cNs is further supported by positive immunoreactivity against Sox2, a stem cell marker, observed in the nuclei of this neuronal population, although at an apparent lower level than in EC (Fig. 6E and F). We next looked for expression of p27Kip1, a marker of cell cycle arrest known to be expressed in neuroblasts and we observed high immunoreactivity in *Rh.* monkey CSF-cN nuclei while it appears lower in EC (Fig. 6G). In contrast, the pattern determinant Pax6 was expressed in Vim positive EC but not in CSF-cNs (Fig. 6H). Altogether, our data indicate that in *Rh.* monkey CSF-cNs do not express maturity markers but rather several proteins classically present in non-mature neurons or neuroblasts.

**Figure 6:**
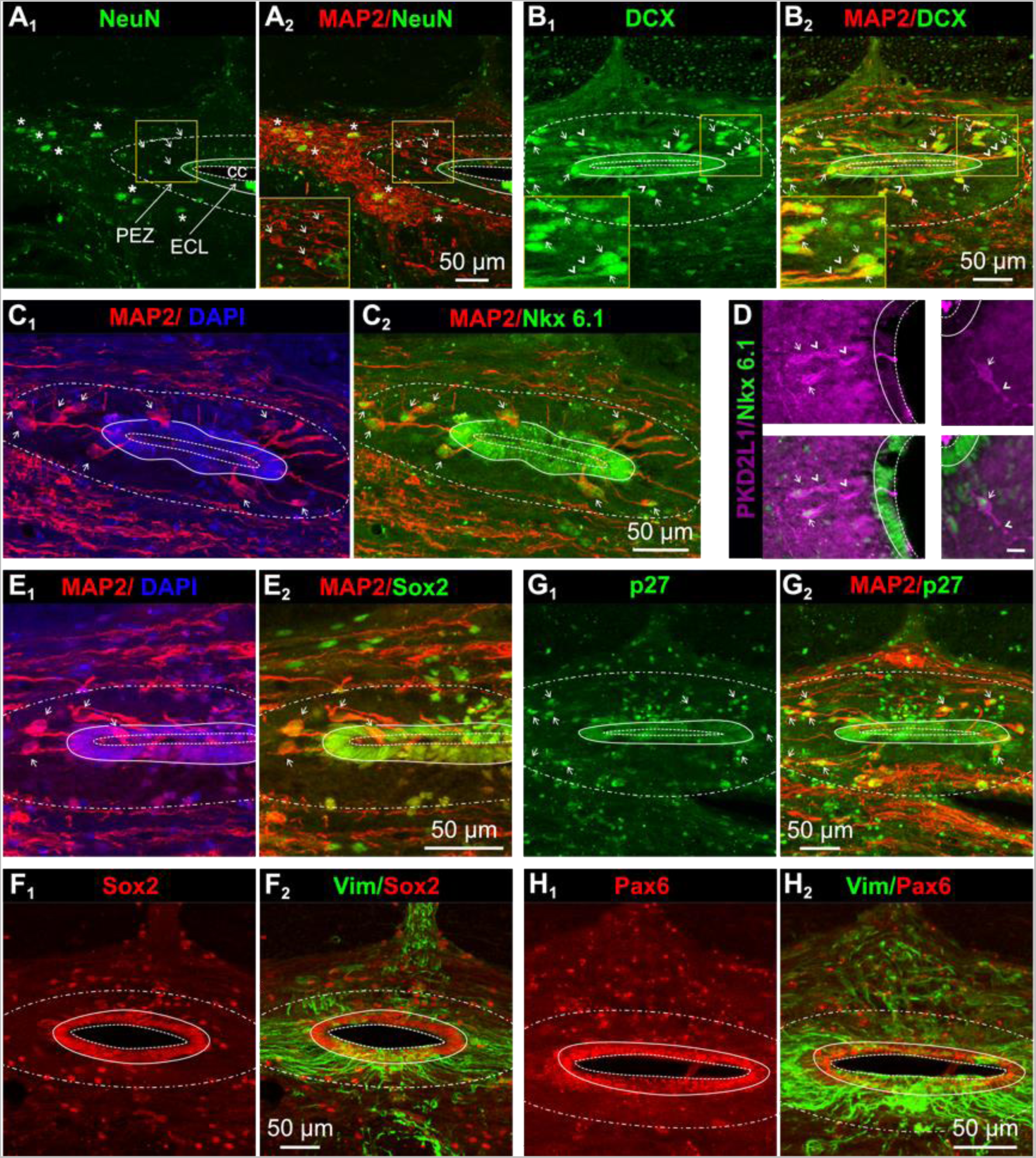
In macaque, CSF-cNs exhibit features of immature neurons. (**A**) Representative micrographs illustrating immunolabelling against NeuN alone (**A1**) and with MAP2 (**A2**) around the cc. Asterisk: NeuN positive neurons in the parenchyma; Arrows: CSF-cNs; Inset in A2: enlargement for the zone around the cc highlighted with a yellow box to illustrate at higher magnification that CSF-cNs are NeuN negative (compare **A1** and **A2**). (**B**) Images illustrating immunolabelling against doublecortine (DCX) alone (**B1**) and with MAP2 (**B2**). Arrows: CSF-cNs; Arrowheads: CSF-cN dendrites; Inset in B1 and 2: enlargement for the zone around the cc highlighted with a yellow box to illustrate at higher magnification DCX/MAP2 co-expression at the somatic and dendritic levels in CSF-cNs (compare B1 and B2). (**C**) Representative micrographs illustrating immunolabelling against MAP2 alone (**C1**) and with Nkx6.1 (**C2**). Arrows: CSF-cNs. Note that Nkx6.1 is expressed by all CSF-cNs and by EC but is absent in the parenchyma. (**D**) Immunolabelling against PKD2L1 alone (magenta, Top row) and with Nkx6.1 (green, Bottom row) in transverse section prepared from cervical SC. The data confirms that all CSF-cNs are PKD2L1+ and selectively express Nkx6.1. (**E**-**F**) Micrographs illustrating the immunolabelling against MAP2 alone (**E1**) or Sox2 alone (**F1**) as well as co-immunolabelling against Sox2 and MAP2 (**E2**) or Sox2 and Vim (**F2**). Note the high Sox2 immunoreactivity of EC around the cc (**E2** and **F1**) as well as Sox2 selective expression in CSF-cNs (Arrows in **E**). (**G**-**H**) Micrographs illustrating the immunolabelling against p27Kip1 alone (**G1**) or Pax6 alone (**H1**) as well as co-immunolabelling against p27Kip1 and MAP2 (**G2**) or Pax6 and Vim (**H2**). All CSF-cNs and EC express p27Kip1 while Pax6 is only present in EC. Note the presence in the PEZ of Pax6 positive cell bodies. Illustrated sections were prepared from SC cervical segments. For abbreviations see Figure 1 and color code for the labels on top of images corresponds to the illustrated markers. Scale bars are shown on the images.

### 5. CSF-cN axonal projection and synaptic contacts

Here, we explored the projection path of putative CSF-cN axon in the SC tissue as well as looked for synaptic contacts on CSF-cNs. Since our data indicate that CSF-cN axon are not labelled with NFH or NF160, two classical axonal markers (see Fig. 1 and 2), we performed our analysis using double immunolabelling against β3-Tub and MAP2. Indeed β3-Tub is an element of cytoskeletal microtubules, present both in the somato-dendritic and axonal compartments, while MAP2 only labels soma and dendrites. This approach, therefore, allows distinguishing the somato-dendritic from the axonal compartments and visualizing axonal projection arising from CSF-cN cell bodies within the PEZ (Fig. 7). Our data on transverse SC sections reveals puncta or patches within the PEZ and adjacent to the ventral ECL that are MAP2 negative but β3-Tub positive and therefore most likely correspond to axons that were cut in the section process (white arrowhead in Fig. 7A_1-2_ and Fig. 4). We next analyzed MAP2/β3Tub co-immunolabelling on longitudinal SC sections across the cc region to confirm this observation and as expected, we could observe MAP2/β3Tub positive CSF-cNs soma and dendrites within the PEZ (open white arrowheads in Fig. 7A_3-4_,) but also MAP2 negative/β3Tub positive rostrocaudal fibers that were either isolated or aggregated as fiber tracts (white arrowheads in Fig. 7A_4_ and see also B-D). To confirm our results, we repeated on transverse and longitudinal sections experiments using double immunolabelling against β3Tub and NFH (Fig. 7B) or NF160 (Fig. 7C). As previously reported, we observed high NF160 (Fig. 7B_2,4_) and NFH (Fig. 7C_2,4_) immunoreactivity mainly outside the PEZ however the β3Tub positive fibers/patches were never NF160 or NFH positive (Fig. 7B_2,4_ or (C_2,4_). Note on the transverse section illustrated in (Figure 7B_2_ occasional NF160 positive elements present in the ventral and dorsal PEZ as well as the dense NF160 positive puncta in the dorsal region of the SC that correspond to ascending or descending axonal projection tracts. Our results, in agreement with the intermediate maturity state reported for *Rh.* monkey CSF-cNs (see above) and the previous studies in rodents (Orts-Del’Immagine et al., 2014, 2017), suggest that their axon do not express mature axonal markers and would project in the longitudinal axis within the PEZ.

**Figure 7:**
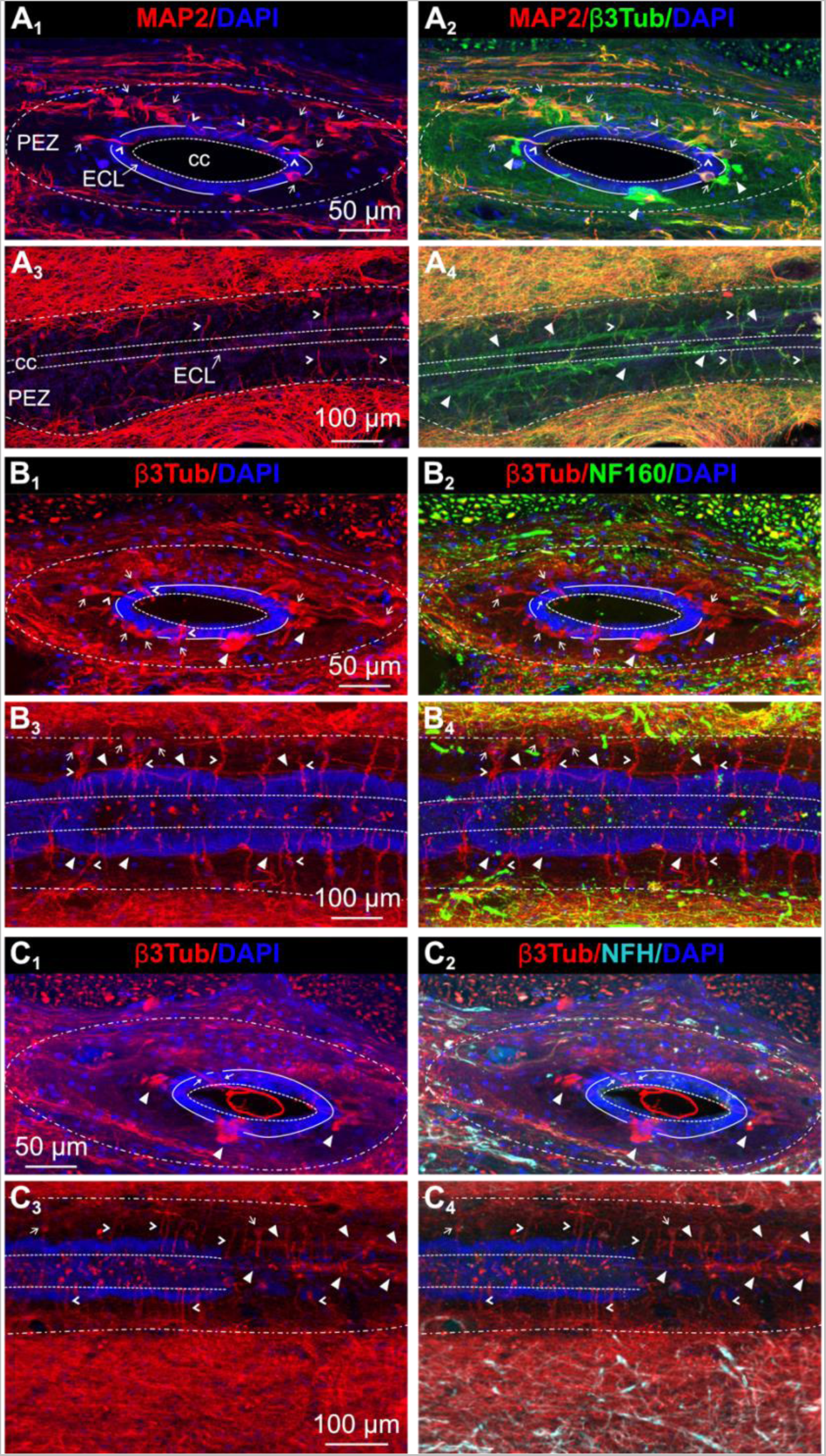
CSF-cN axons appear immature and to project within the PEZ in the longitudinal axis. (**A**) Representative images illustrating the immunolabelling against MAP2 alone (**A1)** and with β3Tub (**A2)** in a transverse or longitudinal (**A3** and **A4**) section. MAP2 labels CSF-cN somato-dendritic compartments while β3Tub also reveals their axonal projection. In **A2**, note in the PEZ the presence of β3Tub positive but MAP2 negative patches (arrowheads) and in **A4**, of β3Tub positive longitudinal fiber bundles (arrowheads) that might correspond to CSF-cN axonal projections. (**B-C**) Representative images illustrating the immunolabelling against β3Tub alone (**B1, B3, C1** and **C3**) and co-immunolabelling against β3Tub and NF160 (**B2** and **B4**) or NFH (**C2** and **C4**) in transverse (**B1, 2** and **C1, 2**) or longitudinal sections (**B3, 4** and **C3,4**). Note in the PEZ, the presence of β3Tub positive patches and longitudinal fiber bundles (arrowheads) that are neither NF160 nor NFH positive. In **C3** and **4**, the section cut with an angle allows visualizing both the CSF-cN cell bodies at the level of the cc (Left) as well as the β3Tub positive longitudinal fiber bundles (Right) at a more dorsal level. For abbreviations see Figure 1. Small arrows: CSF-cN soma; arrowheads: CSF-cN dendrite; filled arrowheads: axonal projection. Color code for the labels on top of images corresponds to the illustrated markers. Scale bars are shown on the images.

Finally, to reveal whether CSF-cNs receive synaptic contacts despite their low maturity state, we analyzed the presence of synaptophysin labelling on MAP2 positive CSF-cNs and explored the phenotype of the identified contacts (Fig. 8). Our results for the cervical segments indicate the presence of synaptophysin positive puncta along CSF-cN dendrites (Fig. 8A) within the PEZ and on their soma but not intra-ependymally. In the thoracic and lumbar regions, most CSF-cNs are intra-ependymal and in these segments, the synaptophysin positive synaptic contacts are mainly observed on CSF-cN cell bodies (Fig. 8B). The orthogonal projections illustrated in (Figures 8A_2,3_ and (B_2,3_ further support this anatomical organization and indicate that the synaptophysin positive contacts form dense immunoreactive puncta along the dendrites and surrounding the whole structure or are present on the soma. To identify the nature of the synaptic contacts CSF-cNs receive, we tested the colocalisation of synaptophysin with markers of cholinergic (ChAT), GABAergic (GAD67), glutamatergic (vGlut1) and serotoninergic (5-HT) neurons. GAD67/synaptophysin double immunolabelling reveals that numerous synaptophysin positive elements contacting CSF-cNs are GABAergic (Fig. 8C_1-3_). Similarly, MAP2/5-HT/ synaptophysin immunolabelling shows that 5-HT fibers are present in the PEZ, mainly in its dorsal part and that they are often colocalized with synaptophysin on CSF-cN soma and dendrites (Fig. 8D_1-3_). In contrast, although we observe ChAT or vGlut1 positive fibers or soma in the parenchyma with some rare fibers arriving in the PEZ, we never visualize on CSF-cNs synaptophysin positive structure that were also immunoreactive for ChAT or vGluT1 (Data not shown). To summarize, CSF-cNs do receive synaptic contacts that are restricted on the somato-dendritic compartment in the PEZ but not within the ECL. These contacts appear to be GABAergic and serotoninergic but not glutamatergic nor cholinergic.

**Figure 8:**
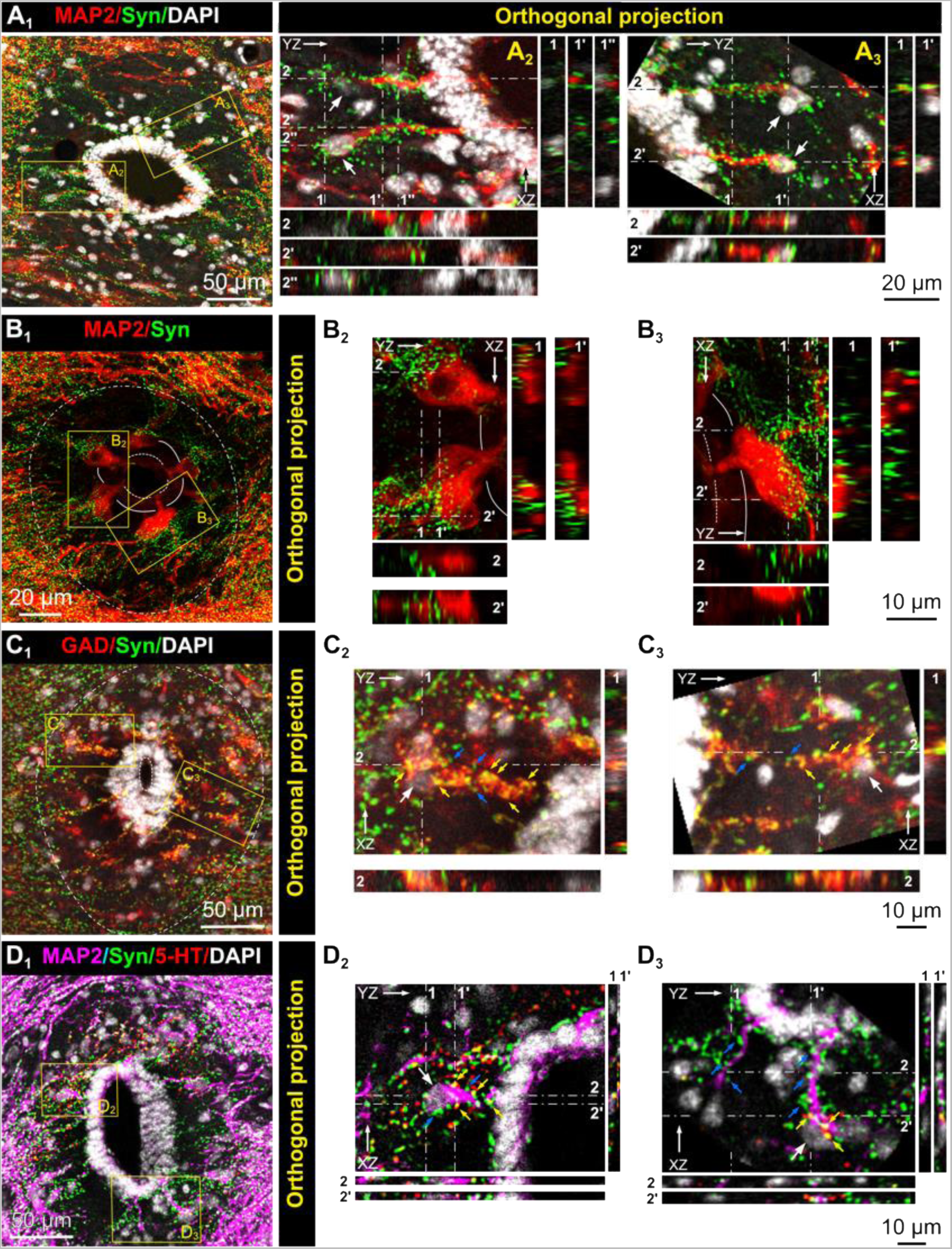
CSF-cN neurons receive somato-dendritic GABAergic and serotoninergic synaptic contacts. (**A**-**B**) Transverse sections showing CSF-cNs (MAP2, red) around the cc and synaptic contacts (synaptophysin, Syn; red) on their soma or dendrites in cervical (**A1)** and thoracic (**B1**) segments (see **A2, 3** and **B2, 3**). (**C**) GAD67 and Syn co-immunolabelling (cervical SC) showing the presence of GABAergic synaptic contacts (**C1**) around CSF-cN dendrites (see **C2** and **3**). (**D**) MAP2, serotonin (5-HT) and Syn co-immunolabelling showing the presence of 5-HT positive synaptic contacts (**D1**) around CSF-cNs cell bodies and dendrites (see **D2, 3**) mainly in the dorsal region of the SC. In **A-D**, Enlargements for the regions highlighted with yellow boxes in **A1-D1** and the respective orthogonal projections (along the YZ axis: 1 and 1’ and the XZ axis: 2 and 2’, see labels). White, blue, and yellow arrows indicate CSF-cN soma, syn positive and GAD (**C**) or 5-HT (**D**) positive puncta, respectively. For abbreviations see Figure 1. Color code for the labels on top of images corresponds to the illustrated markers. DAPI is presented in white. Scale bars are shown on the images.

## DISCUSSION

Our analysis of the cc region in *Rh.* monkey highlights the presence of a large PEZ delimited by radial fibers originating from EC and enriched with astrocytes and microglia. Further, the PEZ is so-called hyponeural with the sole presence of CSF-cNs. This peculiar organization of the region around the cc that extends throughout the entire medullo-spinal axis appears specific to *Rh.* monkey and represent a major difference with that observed in rodents. Nevertheless, in *Rh.* monkey CSF-cNs share with those in rodents a morphology and an immature phenotype in agreement with previous reports. We finally indicate that these neurons receive on their somato-dendritic compartment synaptic contacts from GABAergic and serotoninergic terminals but not cholinergic or glutamatergic ones.

### 1. CSF-cNs specific properties are conserved in *Rh.* monkey

In agreement with previous studies (LaMotte, 1987; Alfaro-Cervello et al., 2014; Djenoune et al., 2014), we show that CSF-cNs are present around the cc in *Rh.* monkey. Although they are only poorly and partially detectable with PKD2L1 antibodies, they are clearly labelled with antibodies against MAP2 and β3Tub. A similarity between rodent and *Rh.* monkey CSF-cNs is also their GABAergic phenotype as indicated by GAD67 immunoreactivity (Orts-Del’Immagine et al., 2014; Djenoune et al., 2017; Jurčić et al., 2021) and morphology. As in mice, CSF-cNs in *Rh.* monkeys have a small ovoid shape and extend a dendrite toward the cc, with a terminal bud in contact with the CSF. Moreover, we show that they are present in *Rh.* monkey along the whole medullospinal axis at various distances from the cc depending on the rostrocaudal level. Thus, CSF-cNs are more distant from the cc at rostral levels (medulla and cervical levels) and closer at thoracic and lumbar levels. Such difference in CSF-cNs distribution along the rostrocaudal and dorsoventral axes has also been described in rodents (Orts-Del’Immagine et al., 2014, 2017) and one could hypothesize that this may be related to a functional heterogeneity and specific projection path as reported in zebrafish (Djenoune et al., 2017). CSF-cNs in the medulla and cervical regions were generally more distant from the cc in *Rh.* monkey than in mice, although a population of distal PKD2L1+ neurons have been recently described in the mouse SC (Tonelli Gombalová et al., 2020; Jurčić et al., 2021) notably in the ventral region.

In *Rh.* monkey alike in rodents, CSF-cNs exhibit properties of classical neurons, illustrated by MAP2 expression, synaptic contacts, neurotransmitter production (GAD67+). Nevertheless, they do not express NeuN, NF160 or NFH classically found in mature neurons but instead we observe expression of DCX, Nkx6.1, Sox2, p27Kip1, all proteins characteristic of neuroblasts or immature neurons. Again, this has previously been shown in rodents (Stoeckel et al., 2003; Sabourin et al., 2009; Kutna et al., 2013; Orts-Del’Immagine et al., 2014, 2017)- and this incomplete maturity profile appears a common feature for CSF-cNs in all analyzed vertebrates. Some characteristics of juvenile neurons have also been observed at the functional level in mice using electrophysiological recordings (Jurčić et al., 2019).

Although it has been shown in rodents that CSF-cNs are not generated at post-natal stage or during adult neurogenesis (Marichal et al., 2009; Kutna et al., 2013), they may conserve some features and functions of juvenile neurons in adult animals. Such unique characteristic for CSF-cNs has been related to their environment in the ependymal stem cell niche and potentially their contact with the CSF. Thus, in *Rh.* monkey CSF-cNs are surrounding the ECL where their dendrites follow the radiating ependymal fibers. Such preferential link between CSF-cNs and EC/EC fibers has also previously been shown in rodents, even for distal PKD2L1+ neurons that are mostly aligned along the ventral ependymal fibers (Jurčić et al., 2021). This unique feature for CSF-cNs, together with their low-maturity phenotype, reinforce the hypothesis for a function as a constitutive element of the stem cell niche along with their chemosensory and mechanosensory function. It further raises the question of whether in mammals they could be activated following SC injury and recruited in regenerative processes. To date this is an unresolved issue and dedicated studies demonstrating CSF-cN activation following SC injury or neuroinflammation need to be conducted.

### 2. A specific astroglial peri-ependymal compartment for CSF-cNs in primates

In rodents, CSF-cNs are mostly located within the ECL or just beneath (subependymal zone) in an area containing astrocytes but also other types of neurons (Djenoune et al., 2014; Orts-Del’Immagine et al., 2014). In contrast, we show that in *Rh.* monkeys CSF-cNs are the only neurons to occupy a specific and wide hyponeuronal PEZ, which is almost completely devoid of other neuronal elements as revealed by the absence of or low immunolabelling against NeuN, NFH and NF160 within this zone as well as MAP2+ staining in the parenchyma at the boundary of the PEZ. Due to this specific anatomical organization for CSF-cNs, MAP2 can be considered as a suitable marker for these neurons in primates since it allows revealing almost the whole population of CSF-cNs distant from 0 to 150 μm to the cc. In addition, this specific PEZ zone also contains a dense synaptic innervation on CSF-cN somato-dendritic compartment (potentially corresponding to the rare NFH or NF160 positive structures observed in the PEZ). This innervation appears mainly GABAergic and could represent at least partially recurrent synaptic contacts between CSF-cNs as recently demonstrated in mice (Gerstmann et al., 2022; Nakamura et al., 2023). Beside this GABAergic innervation, our data indicate that CSF-cNs, mainly in the dorsal PEZ, are also contacted by serotoninergic fibers. Although the origin of these fibers remains to be established, this observation suggests that CSF-cNs might be under the control of supraspinal descending serotoninergic projection. It further raises the question whether CSF-cNs might be contacted by other types of neuromodulatory peptidergic terminals. Concerning CSF-cN axonal projection, our data in *Rh.* Monkeys suggest that they would run within the PEZ in a rostro-caudal and horizontal orientation. This is supported by the presence within the PEZ of longitudinal fibers that were β3Tub+ and MAP2- as well as negative to labelling against NFH/NF160. This organization appears different from that observed in zebrafish, mice or rat, in which axon were shown to project toward the ventral fissure and form long horizontal fiber bundles (Stoeckel et al., 2003; Jurčić et al., 2021; Gerstmann et al., 2022; Nakamura et al., 2023). However, CSF-cN connectivity and the identification of their local and distal neuronal partners is far from being characterized in prrimates and would need to be further addressed using specific tracing approaches. These data would certainly provide insights about their potential function in primates.

Paralleled by the absence of neuronal elements (except CSF-cNs), the *Rh.* monkey PEZ is enriched with astrocytes in agreement with the study from Alfaro-Cervello and colleagues (2014). A high density of astrocytes around the ECL was also observed in mice (Jurčić et al., 2021), suggesting that these cells may play a role in the interface between the CSF and the parenchyma and might interact with CSF-cNs. Moreover, we found that the *Rh.* monkey, the PEZ also contains numerous microglial cells, with some microglial fibers surrounding and even, in some cases, crossing the ECL toward the CSF. Such structural and functional interaction between microglial and EC has recently been shown in mice, notably in the control of EC fate after SC injury (Chevreau et al., 2021). We do not know the functional relevance for such an organization, but we can hypothesize an interaction between astrocytes, microglia and CSF-cNs to control CSF diffusion/composition and regulate the communication between CSF and parenchyma as well as protect the neural tissue against potential toxins or pathogen agents. CSF-cNs might further serve as signal integrator to convey information about the physio-pathological state of the central nervous system and representing a novel interoceptive system enabling the central nervous system to sense itself. This possibility would need to be address in future dedicated studies.

### 3. A phylogenetic reorganization of the peri-ependymal zone?

The different cell organization of the cc environment between rodents and *Rh.* monkey (absence and presence of the PEZ, respectively) raises the question of the anatomical and functional evolution of this PEZ region amongst vertebrates and mammals during evolution. It further questions whether there is difference amongst the primate phylum and whether these data can be translated to Humans. Here we show in *Rh.* a non-human primate (NHP) from the old-world and exhibiting ‘thumb-index’ grip, a specific organization of the PEZ with CSF-cNs as the only neurons present in this region. In contrast, preliminary data obtained in marmoset, a new-world NHP without grip capability and of smaller size, indicate a region around the cc similar to that found in rodents with CSF-cNs within the parenchyma in the ECL or sub-ependymal zone. At this stage we are not able to draw conclusion about the different organization and whether it is related to the phylum, the ability to exhibit the ‘thumb-index’ grip (*eg.* macaque *vs.* marmoset) or to the animal size. To further address this question, it is necessary to extend the characterization of the cc organization and of CSF-cN properties in other primate species.

Concerning Humans and in contrast to other mammalian species, the cc is thought to be mostly occluded and disassembled, at least in adult or old subjects. It is however patent in infants and in the adult medulla and most rostral cervical region (Yasui et al., 1999; Alfaro-Cervello et al., 2014; Garcia-Ovejero et al., 2015; Becker et al., 2018). In the case of occlusion, the cc is replaced by a highly vascularized cell ‘amas’ constituted of bi- and multiciliated ependymal and glial cells (Alfaro-Cervello et al., 2014; Becker et al., 2018). Although, the cell organization of the peri-cc region (putative PEZ) remains to be further characterized in Human, based on electron microscopic (Alfaro-Cervello et al., 2014) and transcriptomic studies (Ghazale et al., 2019), CSF-cNs have not yet been detected in Humans even if β3Tub immunoreactive processes have been observed in the cc region (Garcia-Ovejero et al., 2015; Becker et al., 2018; Torrillas de la Cal et al., 2021). To data there is no definitive data to confirm or infirm the presence of CSF-cNs and further immunohistochemical investigations using specific neuronal markers will be necessary in Human SC and medulla samples, notably with a patent cc, to putatively identify CSF-cNs and specify the PEZ organization relative to NPH and rodents. On the other hand, if CSF-cNs are not present in Human although they have been observed in many vertebrate species and in NHP, it will be interesting to consider why and when they would have disappeared during evolution and by which cellular type, if any, they were replaced.

## CONCLUSION

Here, we report an in-depth anatomical study for the region around cc in the *Rh.* monkey that appears to be also present in Human. Although highly challenging in NHP, it will be necessary in the future to pursue the characterization of the PEZ and the CSF-cN properties, identify their axonal projection and post-synaptic partners to come closer to the understanding of their function. Finally, the functional ground for this unique organization of the PEZ in *Rh.* monkey compared to rodents is unclear and would need to be further characterized. One could hypothesize that it represents a buffer zone to control diffusion of active molecules from the CSF towards the parenchyma with CSF-cNs as intrinsic sensory neurons that would serve to sense signals and convey the information to regulate the central nervous system homeostatic state.

## MATERIAL & METHODS

### Animals

Experiments were conducted on 5 adult *Macaca mulatta* (*Rh.* monkey; weight between 2 and 7 Kg) that have been used in experimental protocols carried out in the institute for other projects. To comply with the 3Rs rules, they were made available at the end of these procedures to collect medullo-spinal tissues. Housing condition and all experiments were conducted in conformity with the rules set by the EC Council Directive (2010/63/UE) and the French “*Direction Départementale de la Protection des Populations des Bouches-du-Rhône* (DDPP13)”. Anesthesia was induced with an injection of zoletil (10 mg.Kg^-1^), xylazine (1 mg.Kg^-1^) and buprenorphine (0.03 mg.Kg^-1^). Following full reflexes loss, an intravenous injection of pentobarbital (100 mg.Kg^-1^) was performed. Upon confirmation of respiratory and cardiac arrest, transcardiac perfusion was carried out. For immunohistology on rodent tissues, 6-week-old mice and 3 month-old rats (3 animals for each species) were injected 30 minutes prior to the procedure with the non-steroidal anti-inflammatory meloxicam (5 mg.Kg^-1^) and subsequently anesthetized with intraperitoneal administration of 100 mg.Kg^-1^ ketamine (Carros, France) and 10 mg.Kg^-1^ xylazine (Puteau, France). Before skin incisions, the local antalgic lidocaïne (5 mg.Kg^-1^) was injected subcutaneously. Animals were transcardially perfused initially with 0.1 M PBS followed by 4% paraformaldehyde (PFA; Electron Microscopy Sciences) in 0.1 M PBS solution. Tissues were immediately collected, separated in the segments of interest (medulla, cervical, thoracic, lumbar, and sacral levels), post-fixed in PBS-4% PFA at 4°C overnight, cryoprotected for several days in 30% sucrose at 4°C and finally frozen in isopentane (−40°C) after OCT embeding.

### Immunohistochemistry

Transverse and longitudinal thin sections (30 μm) were prepared using a cryostat (Leika CM3050) and collected serially in 24-well plates containing 0.1 M PBS. For further conservation, sections were transferred in a cryoprotecting solution at −20°C until use. For immunohistochemistry, sections were washed in PBS, incubated first 1h in PBS containing 0.3% Triton X-100 (Sigma) and then in a blocking solution containing 3% horse or goat normal serum and 1% bovine serum albumin (BSA). Sections were incubated overnight at 4°C with primary antibodies (see list in Table 2) in an incubation medium (PBS with 3% horse or goat normal serum). Sections were then washed at least 3 times in PBS and incubated for 2h with secondary antibodies conjugated to either AlexaFluor 488, 594 or 647 (Life Technologies) and finally washed in PBS several times. When more than one primary antibody was used in an immunohistofluorescence labeling experiments, the primary antibodies were applied sequentially. Sections were mounted on polylysine coated slides and coverslipped with a mounting medium for fluorescence microscope preparation (generally with DAPI staining). To assess the selectively of the observed immunolabelling, in each set of experiments, some sections were treated in the absence of primary antibodies and using secondary antibodies alone (negative control). Each experiment was performed on tissues obtained from 2-3 animals and analyzed on 3-6 sections for each segment of interest. Most antibodies (NeuN, GFAP, Vim, β3Tub, DCX, Nkx6.1, MAP2, PKD2L1) have been previously used in our studies in mice (Orts-Del’Immagine et al., 2014; Jurčić et al., 2021) and were validated in *Rh.* monkey by the labelling observed in similar cell populations as in rodent as well as positive labelling for NeuN, MAP2, ChAT, β3Tub, NFH, NF160 and synaptophysin in neurons and neuronal elements present in the same section away from the cc. Vim, GFAP and Iba1 immunofluorescence was detected as expected in EC, astrocytes, or microglial cells, respectively. The antibodies Sox2, Pax6, p27Kip1 were validated by the expected immunolabelling of the cc ependymal stem cell population.

**Table 2:** List of primary antibodies used for immunolabelling.

| **Antibodies** | **Distributor** | **Reference** | **Host** | **Dilution** |
| --- | --- | --- | --- | --- |
| **β3-tubuline/**<br>**TUJ1** | Sigma | T2200 | rabbit | 1:500 |
| **β3-tubuline**<br>**TUJ1** | Biologend | 801213 | mouse | 1:500 |
| **ChAT** | Millipore | AB144P | goat | 1:100 |
| **DCX** | Santa Cruz | SC-8066 | goat | 1:100 |
| **GAD67** | Millipore | MAB5406 | mouse | 1:500 |
| **GFAP** | Sigma | G3893 | mouse | 1:500 |
| **GFAP** | Dako | Z0334 | rabbit | 1:500 |
| **5-HT** | Immunostar | 20079 | goat | 1:500 |
| **Iba1** | Wako | 019-19741 | rabbit | 1:800 |
| **MAP2** | Sigma | M-1406 | mouse | 1:600 |
| **NeuN** | Millipore | MAB377 | mouse | 1:800 |
| **NF160** | Sigma | N5264 | mouse | 1:1000 |
| **NFH SMI32** | Biologend | 801702 | mouse | 1:500 |
| **Nkx6.1** | DSHB | F55A10 | mouse | 1:100 |
| **P27Kip1** | Epitomics | 2187 | rabbit | 1:200 |
| **Pax6** | Santa Cruz | Sc-53108 | mouse | 1:100 |
| **PKD2L1** | Millipore | AB9084 | rabbit | 1:700 |
| **Sox2** | Santa Cruz | Sc-365823 | mouse | 1:200 |
| **Synaptophysin** | Epitomics | 1485 | rabbit | 1:500 |
| **V-Glut** | Millipore | ABN1647 | rabbit | 1:200 |
| **Vimentin** | Millipore | AB5733 | chicken | 1:700 |

### Image acquisition and analysis

Sections were observed on a confocal laser scanning microscope (CLSM; Zeiss LSM700) equipped with solid state fiber optic lasers and single plane images or stacks of images were acquired with a 0.5-1x digital zoom using a 20x objective (numerical aperture, NA: 0.8 for an optical thickness of ∼2 µm). Images were acquired at optimal resolution of 1024×1024 pixels. When using 2-4 fluorochromes with different excitation/emission spectra, images were acquired sequentially for each channel (488 nm, green; 555 nm, red; 405 nm, blue; 639 nm, magenta) and the filters and photomultiplier tubes settings chosen to optimize images and avoid signal crosstalk. Images were analyzed and prepared using ZEN 2009 light Edition (Zeiss software), ImageJ 1.45 (NIH) and Corel Photo-Paint. For better visualization, contrast and brightness were adjusted in images used for the figures. Orthogonal projection and co-expression analysis were performed in ImageJ using dedicated plugins. Images illustrated at low magnification were obtained using the ZEN 2009 mosaic routine (from 4 x 4 to 6 x 6 fields of view) while high magnification images were obtained from z-projection of 5 to 12 images in a stack. For quantification analysis of the cell density, MAP2+ CSF-cNs were counted manually in each image and the total cell number divided by the section thickness (10 µm stack depth). For the analysis of PEZ width, cc area (10^3^ x µm^2^), CSF-cN distance to the cc, the measures were performed using ImageJ.

### Statistical analysis

All data presented in (Figure 5 are expressed as mean ± SD and represented as whisker boxplots using the Tukey’s method as well as with “violin” plots to illustrate data point density and the data point distribution. In whisker plots, for each data set, the median and the 25th to the 75th percentiles (lower and upper limits of each bar, respectively) are calculated. Next, the interquartile distance (IQR) is determined as the difference between the 25th and 75th percentiles, and the whiskers limits, or “inner fences” calculated as the 75th percentile plus 1.5 times IQR and the 25th percentile minus 1.5 times IQR. We compared one parameter (PEZ width, CC area, CSF-cN number or distance to the cc) across several medullo-spinal segments, here labeled as *Region*. In one animal, the parameters measured in each *Region* are assumed dependent from the organization of the other regions, and the variable *Region* represents a dependent factor. On the other hand, the same variables represent independent factors when considered in different animals. Therefore, the statistical analyses required using a nonlinear mixed effect (nlme) model taking into account in a hierarchical way both dependent and independent variables as well as the interaction between the variables. Finally, the choice of animals used for this study was done randomly among all those available; therefore, it is necessary to implement the statistical model with a factor corresponding to this random effect. The statistical analyses were carried out using the R Studio 3.3.1 statistical software (R Studio Team, 2015) in the R environment and the “non-linear mixed effect (nlme version 3.1–128)” package (Pinheiro et al., 2004). The null hypothesis was set as the absence of difference between the data among each group. The nlme model was tested using an analysis of the variance (ANOVA), and a subsequent post-hoc Sidak test for multiple pairwise comparison. We provide the values for χ^2^, the degree of freedom (number of parameters −1, dF) and the p-values for the given fixed factor tested. Statistical differences were considered as significant for p < 0.05.

## ADDITIONAL INFORMATION

### Data availability statement

The data of this manuscript are available upon request.

### Authors contributions

All experiments were performed at the ‘Institut des Neurosciences de la Timone (INT)’ of Aix-Marseille University. Conception and design of the work: AK & NW. Acquisition, analysis, and interpretation of data for the work: AK & NW. Drafting the work or revising it critically for important intellectual content: AK & NW. All authors have read and approved the final version of this manuscript and agree to be accountable for all aspects of the work in ensuring that questions related to the accuracy or integrity of any part of the work are appropriately investigated and resolved. All persons designated as authors qualify for authorship, and all those who qualify for authorship are listed.

## Acknowledgments

We gratefully thank Caroline Blanc-Tailleur and Jorge Jose Franco-Ramirez for their assistance in immunohistochemistry experiments and analysis, Edith Blasco for her help in the quantitative analyse as well as SpiCCI team members for careful proof reading and comments. We also thank Hugo Boulanger, Joao Xavier Cardoso and Alexandre Laine for their contribution during their intership. We acknowledge the *Institut de Neurosciences de la Timone* (INT) technical facilities (NeuroBioTools/ConnectoVir: Molecular Biology and Histology; INPHIM: confocal microscopy) for their support in the study and in particular Jean-Alban Rathelot, Luc Renaud, and Eduardo Gascon for their help in primate tissues extraction. We acknowledge the Mediterranean Primate Research Centre (MPRC) for providing NHP animals.

